# Exploitative competition among different dormancy strategists shapes community diversity, recovery from perturbations and response to dispersal

**DOI:** 10.64898/2026.11.18.767651

**Authors:** Andreas Ulmer, Ashley Shade, Rosalind J Allen

## Abstract

Dormancy is a reversible physiological state of reduced metabolic activity that occurs in plants and animals, but is particularly common among microorganisms. Microorganisms can switch into dormancy spontaneously or in response to environmental cues. While previous work has investigated the ecological and evolutionary consequences of the switching mechanism, it is unknown how different dormancy-switching mechanisms within a microbial community interact to shape composition and community-level response to environmental changes.

Here, we used mathematical modelling to investigate how different microbial strategies for entering and existing from dormancy shape community dynamics under stable conditions, following perturbation, and after dispersal. We modelled strategists that had no dormant state (non-switching), those that employed spontaneous switching, those that switched given a cue (triggered switching), and those that used a combination of spontaneous and triggered switching (mixed switching). We also examined how competition and the costs of switching influence activity dynamics within the community.

Our simulations revealed the importance of exploitative competition among dormancy strategists in shaping the community response to perturbation and dispersal. In a mixed community containing different dormancy strategists, spontaneous switchers maintained a dormant pool, but suffered an implicit competitive disadvantage. Meanwhile, triggered switchers were more competitive under favourable conditions but could be pushed into dormancy by exploitative competition from other strategists. Interestingly, dispersal events could intensify competition, induce dormancy in resident populations and alter the relative activity of different strategists, while also restoring community members lost during perturbation.

To evaluate these predictions, we then analysed an existing dataset of soil mesocosms that experienced heat perturbation and subsequent dispersal. The data showed a stable dormant pool, consistent with the presence of stochastic switchers, and the observed dormancy dynamics following dispersal supported competition-driven inactivation among triggered switching strategists. However, we found no evidence that dormancy provides general protection against the heat perturbation. Instead, the results suggest that survival may depend on the inherent tolerance of community members, while subsequent recovery reflects the interplay between dormancy, immigration, and competition.

Our work shows how dormancy not only impacts community-level resilience to perturbations but also alters, and is altered by, exploitative competition among community members.

## Introduction

Dormancy, or the reversible entry into a state of reduced metabolic activity, is a widespread ecological trait, occurring in plants (for example, seeds), animals (hibernation) and microorganisms. The existence of dormant metabolic states and the mechanisms for entering and exiting from them have important ecological and evolutionary implications at the level of individuals, populations and communities(Bradley, 2025; Lennon and Jones, 2011; Letten et al., 2024; McDonald et al., 2024). While dormancy mechanisms in plants and animals have been, in general, well-studied, for microorganisms the diversity of dormancy strategies and their roles for community ecology and evolution remain to a large extent unknown. However, for some microbial communities, a substantial fraction of community members may be inactive at any given time (Guigard et al., 2026; Jones and Lennon, 2010; Pedros-Alió, 2006); indeed, in soil communities, the majority of detected microorganism and taxa can be dormant (Lennon and Jones, 2011), with potential to reactivate and contribute to community responses to perturbation (Barnett and Shade Ecology Letters)(Bandopadhyay et al., 2024). This is especially important since microbial activity underpins many of the Earth’s biogeochemical cycles (Falkowski et al., 2008; Nayfach et al., 2021), the functions of managed and natural ecosystems (Averill et al., 2022; Fierer, 2017; Madsen, 2005; Philippot et al., 2024; Shade et al., 2012) and plant, human and other animal health.

Cellular switching into (and out of) dormancy can in general be classified as spontaneous (Balaban et al., 2004; Sturm and Dworkin, 2015) or triggered. Triggered switching is responsive to environmental factors such as temperature stress (Barnett and Shade, 2024), salt stress (Rath and Rousk, 2015), electrochemical potential (Kikuchi et al., 2022), or interactions with surrounding cells (Dworkin and Shah, 2010; Mutlu et al., 2018). In some cases, both triggered and spontaneous switching can occur in the same population but under different contexts. For example, in *Escherichia coli*, resource depletion can induce a regulated growth arrest that maintains low cellular activity and allows rapid recovery when resources become available (Rotem et al., 2026). By contrast, abrupt starvation can produce a disrupted growth arrest with larger recovery times (Rotem et al., 2026).

Theoretical studies have suggested that spontaneous dormancy may function as a bet-hedging mechanism that helps a population to survive unpredictable environmental fluctuations (Jones and Lennon, 2010). It has also been suggested that spontaneously awakening “scout” cells could signal back to kin to confirm a favourable environment (Buerger et al., 2012; Epstein, 2009). Other studies have considered the possible evolutionary benefits of spontaneous vs triggered switching strategies, with the concept that spontaneous vs triggered switching strategies present a trade-off in precision of responsiveness to environmental changes versus energetic investment (Karakoç et al., 2026). Which strategy is optimal may depend on the characteristics and dynamical fluctuations of the environment (Kussell et al., 2005; Lastad and Mitarai, 2025). In a complementary view, it has been proposed that triggered switching into dormancy may decrease competition between cells (Gardner et al., 2007) with some likely competitors of *Bacillus* being able to induce its germination by manipulating amino acid cues in the environment (Comerci et al., 2026; Mutlu et al., 2018). From the perspective of a microbial community, either triggered or spontaneous switching is expected to give rise to subpopulations of dormant microorganisms; this dormant pool can be viewed as a “seed bank” that may survive environmental changes and subsequently resuscitate when conditions become favourable (Lennon and Jones, 2011).

Competition is another important ecological process that shapes population dynamics, community assembly and the maintenance of diversity (HilleRisLambers et al., 2012) (Chesson, 2000; Letten et al., 2017). In microbial communities, competition occurs both through limitation of shared resources (exploitative competition) and through direct inhibition of competitors (interference competition) (Hibbing et al., 2010; Ghoul and Mitri, 2016). Exploitative competition is especially relevant for understanding the role of dormancy in microbial community dynamics. Although previous work has proposed that dormant populations that do not consume resources (Gardner et al., 2007) can influence competitive dynamics, this concept has not yet been fully investigated.

Ecological communities function within and are shaped by the dynamic environments. For microbial communities, two aspects are especially important: perturbations and dispersal events (Custer et al., 2022; Evans et al., 2020; Sorensen and Shade, 2020). Perturbations are external events, such as abrupt changes in abiotic conditions, that disrupt communities’ ecological structure (Turner, 2010), for example by causing the local extinction of some community members. Dispersal is the movement of microbes between different environmental locations. Since microbial communities exhibit spatial variation (Fierer and Jackson, 2006; Martiny et al., 2023), dispersal enables microorganisms to colonise new habitats (Mallon et al., 2015). After dispersal, microbial immigration into the resident community can alter its composition and function due to community coalescence (Amor et al., 2020; Custer et al., 2024; Litchman, 2010; Vila et al., 2019). Crucially, dispersal and perturbation are linked ecologically, since dispersal can enable recovery of sensitive populations after a disturbance (Nemergut et al., 2013; Shade, 2023). Therefore the role of dormancy within microbial communities needs to be considered in the context of the interplay between, especially, exploitative competition, perturbation and dispersal events.

The role of activity transitions in microbial community dynamics is in general underexplored, with most studies focusing on individual cells or single-species populations. The implications of activity transitions for understanding community response to perturbation and dispersal are likely significant but have not been fully investigated (Philippot et al., 2021; Shade et al., 2012). As present, it is not only unclear which dormancy switching strategies (e.g. spontaneous vs triggered) occur in which microbial populations or if particular strategies are favoured within particular communities; we also do not know how the presence of different switching strategies within a community influences its responses to perturbation and dispersal. Spontaneous switching may promote survival under unpredictable or unfavourable conditions by maintaining a reservoir of cells that reduces the risk of population extinction after a perturbation. However, triggered switching may also play an important role following perturbation, as disturbances can create newly available niches that can be rapidly occupied by previously dormant microbes that can sense and respond to the environmental change. Dispersal may also have a complex interplay with dormancy switching, since interactions with the newly introduced microorganisms (e.g. niche competition) could trigger dormancy transitions, as well as altering the favourability of dormant vs active states.

As a first step towards addressing these questions, a previous study investigated the dynamics of microbial activity in soil mesocosms subjected to a strong heat perturbation, and then allowed to recover over a study duration of 45 weeks (Sorensen and Shade, 2020). In addition to the heated versus ambient treatments, a subset of the heated mesocosms also received an artificial dispersal event consisting of immigration from a control mesocosm. Significant changes in community composition and in taxon-level activity patterns occurred upon warming, with the community becoming dominated by thermotolerant taxa that remained abundant after perturbation in the absence of dispersal. In addition, dispersal contributed substantially to community resilience (ability to recover post-perturbation): post-perturbation, immigrants from an unperturbed community were able to displace thermotolerant taxa and substantially restore the original community composition (Sorensen and Shade, 2020). Interestingly, taxa were classified according to their characteristic patterns of activity before, during and after the heat perturbation and dispersal events. This work suggests that microbial traits like tolerance to perturbation and ability to immigrate post-dispersal as well as activity switching strategies (e.g. spontaneous vs triggered switching) may interact to drive community-level responses; however the mechanisms by which these different traits interact within a community remain unknown.

Although inactive cells are intrinsically difficult to study via classical microbiological methods, substantial efforts have been made to develop reliable methods to define and track activity states within microbial communities (Blagodatskaya and Kuzyakov, 2013). For example, the ratio of abundances of 16S rRNA versus the 16S rRNA gene can be used as a proxy for the activity state of a given taxon within the community (Blazewicz et al., 2013; Felske et al., 1998; Griffiths et al., 2000; Sorensen and Shade, 2020). Complementary microscopy-based methods can provide information on activity levels of individual microbial cells (Gonzalo et al., 2026). These methods, and others (e.g., Guigard et al. 2026), can be used to differentiate between active and inactive bacteria.

Mathematical modelling is an important tool that can help to understand complex observations in microbial ecology. In a mathematical model, a set of underlying concepts (or “rules”) is postulated and the implications of these concepts for community dynamics is explored and compared with experimental observations (Bäcker et al., 2026; Gardner et al., 2007; Gonze et al., 2018; Letten et al., 2024; Stolpovsky et al., 2011; Widder et al., 2016). For example, niche-based models focus on the concept of an environmental niche in which limited resources may lead to exploitative competition (Ghoul and Mitri, 2016; Kivlin et al., 2021; Malard and Guisan, 2023). This in turn can influence microbial activity strategies, e.g. switching to dormancy could be favoured if dormant cells do not compete for resources (Manzoni et al., 2012; Zhao et al., 2026). A mathematical model complements wet-lab data collection and analysis because it can inform the understanding of the possible underlying ecological principles.

In this work, we develop a niche-based mathematical model to explore how spontaneous and triggered activity switching strategies influence microbial community response to perturbations and the role of dispersal in community recovery after perturbation and then validate the predictions with empirical microbiome data from a soil mesocosm experiment (Sorensen et al. (2020)). We model a microbial community that is exposed to an intense perturbation, followed by a dispersal event. We use our model to test how spontaneous versus triggered switching into and out of dormancy influence microbial coexistence within the niche and recovery of the community from the perturbation. Simulations of our model show how the presence of different strategists within the community varies the dormant pool size and its responsiveness to environmental change. Our simulations reveal a key role for exploitative competition between dormancy strategists: spontaneous switchers are less competitive than triggered switchers in the absence of perturbation but triggered switchers can avoid competition by building up a large dormant subpopulations. Our simulations also show how competition caused by dispersal can induce dormancy in resident populations. Guided by these model-informed hypotheses, we analyzed the dataset of Sorensen et al. (2020) and found that the data is generally consistent with the model predictions; however, in contrast to the expectations of our model and previous models, we do not find evidence that dormant microbes show increased tolerance to the strong heat perturbation applied in that work. Our work shows how a niche-based conceptual model can help to interpret complex dynamical data in microbial ecology and suggests that the interplay between activity switching and competition for niche resources may help shape community-level responses to perturbations and dispersal.

### A niche-based model for microbial dormancy switching

In our model, microbes proliferate and die within a niche. We model the dynamics of four microbial types (or “strategists”), that are defined by the nature of their switching between active and inactive states (Fig. 1A). Non-switching strategists (NSS) do not switch, i.e. they remain active. Spontaneous switching strategists (SSS) switch into and out of dormancy at a rate that is independent of external factors (and is the same in both directions to reduce the number of parameters in the model). Triggered switching strategists (TSS) switch to dormancy in response to resource competition (Gardner et al., 2007). The rate of triggered switching is assumed to be a function of niche occupancy: as more cells occupy a niche, the rate of switching into dormancy increases, whereas the rate of reactivation decreases. The model therefore incorporates a dynamic, or gradual, form of triggered switching, in which the chance of a cell switching increases with changing environmental conditions, rather than an immediate binary switch in which all cells switch when an environmental threshold is reached. Gradual triggered switching might occur, for example, if the trigger were to be spatially distributed and therefore not act simultaneously on all cells. However, by tuning the triggered switching rate parameter (s_TD_, defined later), the model can be tuned between dynamic and binary switching. Mixed switching strategists (MSS) have both spontaneous and triggered switching mechanisms; i.e. they switch to dormancy both spontaneously and in response to competition. This reflects that concept that both spontaneous and triggered switching mechanisms could, in principle, coexist within the same cell. For example, the MSS strategist could represent a ‘scout’ (Epstein, 2009) which awakes randomly, but also responds to a cue which is in our model the niche occupancy, although our model does not incorporate a mechanism by which reactivated scout cells induce the reactivation of other dormant cells.

**Figure 1:**
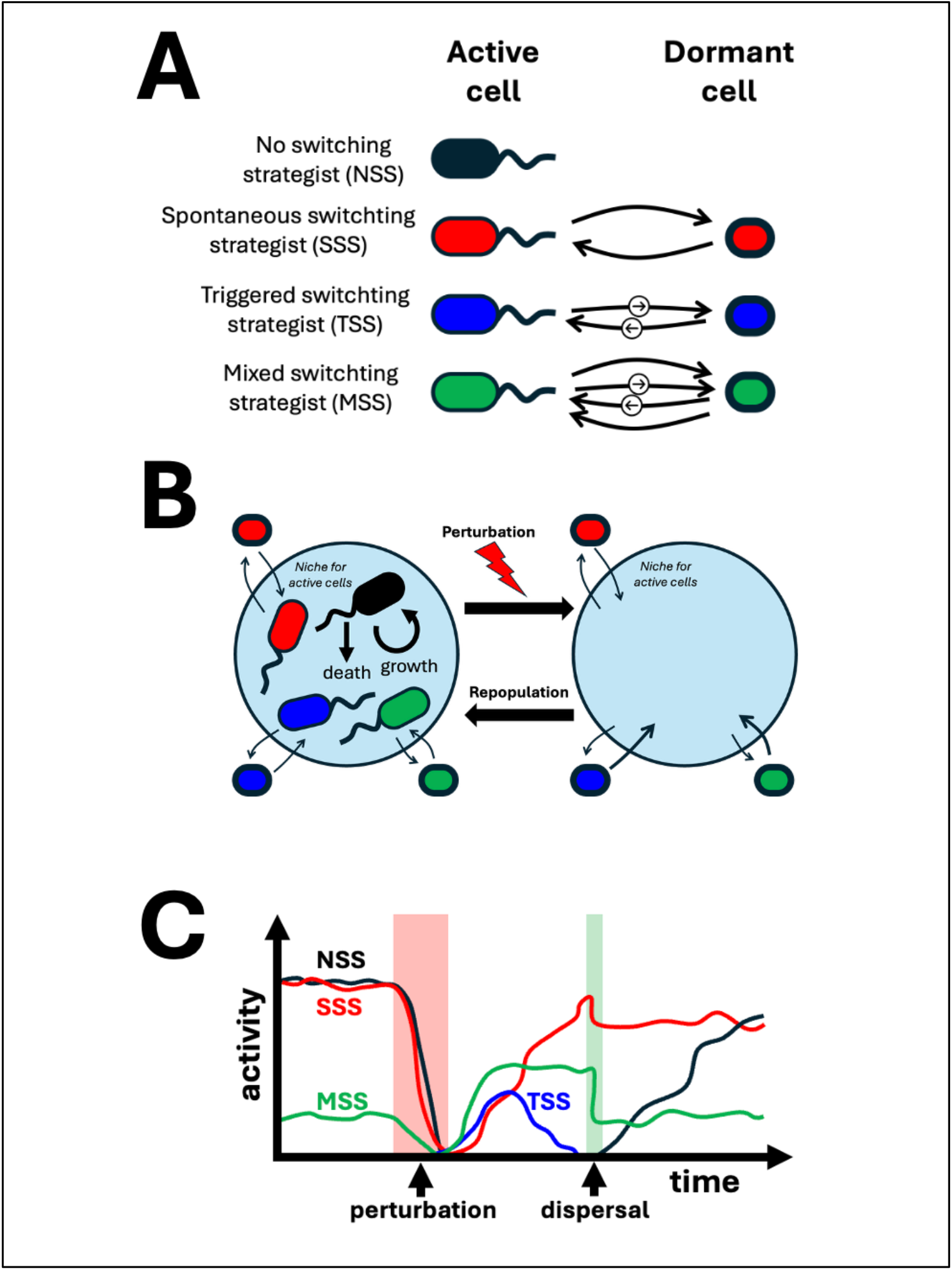
Model concept. **(A)** The model considers 4 dormancy switching strategies. The no switching strategist (NSS; black) remains active, i.e. does not switch to dormancy. The spontaneous switching strategist (SSS; red) switches in and out of dormant state at a fixed rate, independent of the environment. The triggered switching strategist (TSS; blue) switches in and out of dormant state in response to competition for niche capacity. The mixed switching strategist (MSS; green) switches to dormancy both spontaneously and in response to competition. **(B)** Processes within the model. Active microbes proliferate and die within the niche; proliferation stops when the carrying capacity of the niche is reached. Microbes also switch between active and dormant states. Dormant microbes have increased tolerance to the perturbation and do not compete for niche capacity. The colours represent different switching strategists as in A. A perturbation removes all active cells. After perturbation the surviving dormant cells can reactivate and repopulate the empty niche. **(C)** Sketch showing hypothetical responses of different switching strategists to perturbation and dispersal. The NSS is eliminated by the perturbation and only restored by a dispersal event. The SSS survives the perturbation due to its dormant subpopulation and can regrow after the perturbation. It experiences increased competition after the dispersal event. The TSS also survives the perturbation and subsequently expands into the vacant niche, until the recovering SSS and MSS populations displace the active TSS. The MSS shows intermediate dynamics between SSS and TSS.

To account for competition within the niche, the model includes a niche capacity C that limits the total active population size. As the active population size increases, the proliferation rate of the active microbes decreases. This reflects limited availability of resources; it is assumed that all active microbes compete for the same resource pool. It is assumed that dormant cells do not consume resources and hence the number of dormant cells is not limited in the model. As explained above, for the triggered switching strategists (TSS and MSS), we assume that resource competition triggers switching to dormancy, i.e. the switching rate is coupled with the difference between the total active population and the niche capacity C.

To represent an intense perturbation in a simple way, in our model a perturbation is assumed to instantly kill active but not dormant cells (i.e. all active microbes are removed from the system at a specified time point, while leaving the number of dormant microbes unchanged). This reflects the fact that some inactive cellular states, such as spores, are known to provide increased tolerance to environmental perturbations (Setlow, 2014a).

A dispersal event is modelled by introducing immigrant microbes into the system. We model this process by adding a predefined community composed of a mixture of active and dormant cells (Fig. S2) at a specified time point. The added community corresponds to a simulated unperturbed community with the same parameter values (see Methods).

We use a deterministic, Lotka–Volterra-style modelling framework (Altermatt et al., 2011; Gonze et al., 2018) in which we track the sizes of the subpopulations of active and dormant microbes for each strategist as a function of time. For the non-switching strategist (NSS), there is no dormant subpopulation so the dynamics of this strategist is described by a single ordinary differential equation:

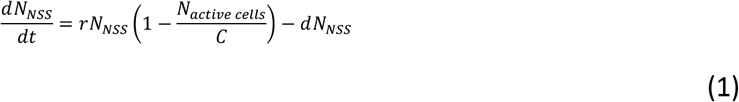

Here, *N_NSS_* represents the number of (active) NSS microbes present at time t, *N*_active cells_ is the total number of active microbes in the system (of any type) and C, r and d are parameters describing the carrying capacity of the niche, the maximal growth rate and the intrinsic death rate, respectively. The first term in Eq. (1) describes growth of the NSS microbes that occurs exponentially at rate r when the niche is empty but slows down as the number of active microbes in the niche approaches the carrying capacity C. The second term describes death of the NSS microbes, which occurs spontaneously at a fixed rate d.

The dynamics of the spontaneous switching strategist (SSS) is described by two equations corresponding to the active and dormant subpopulations, *N_SSS_*_,active_ and *N_SSS_*_,dormant_ respectively:

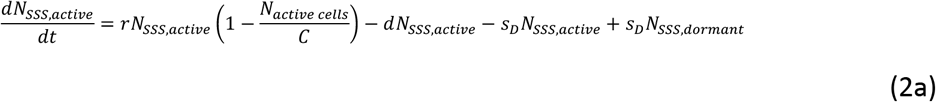

and

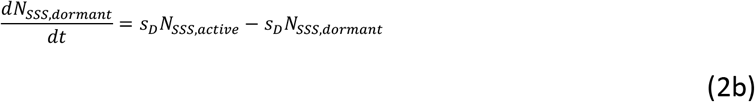

Eq. (2a) accounts for proliferation and death of active cells (as in Eq. (1) for NSS) and also contains terms that describe spontaneous switching from activity to dormancy and vice versa. It is assumed that the switching rates in either direction are equal (the switching rate is here denoted s_D_). Eq. (2b) expresses the fact that the dormant subpopulation switches reversibly between dormant and active states but neither proliferates nor dies.

The triggered switching strategist (TSS) is described by similar equations to those for SSS, but with different rates for switching into and out of dormancy:

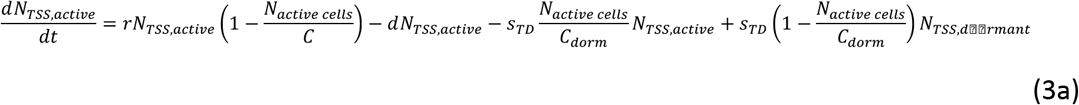

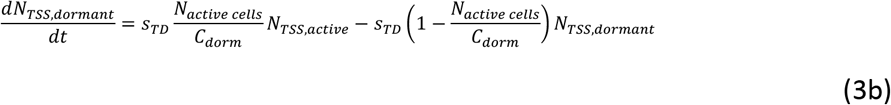

In Eqs. (3a) and (3b), the switching from activity to dormancy depends on the ratio N_active cells_/C_dorm_. Here, *C*_dorm_ = *C*(1 − *d*) is the number of active cells that are present when the niche is fully occupied (this is similar to the niche capacity C but is slightly reduced because of cellular death) and s_TD_ is the switching rate. The rate of switching into dormancy is low when competition is low (i.e. only few active cells are present) but it becomes higher when competition increases (i.e. the number of active cells approaches the maximum possible number within the niche). The rate of reactivation, i.e. switching from dormancy to activity, is also scaled, by a factor 1 − *N*_active cells_/*C*_dorm_, to ensure that the reactivation rate becomes zero when the niche is fully occupied. The triggered switching rate parameter s_TD_ can be tuned to model dynamic triggered switching, in which the population response gradually to a change in environmental conditions (low value of s_TD_), versus binary triggered switching, in which the whole population responds rapidly to the change (high value of s_TD_).

To account for the possible cost of sensing local competition, in some simulations we reduced the growth rate parameter r (Bruggeman et al., 2020) by a factor of 2 for the TSS and MSS strategists compared with the value for the NSS and SSS strategists. This assumption is based on resource allocation considerations, since sensing mechanisms require proteins, enzymes, or other cellular machinery which cost e.g. energy (Govern and Ten Wolde, 2014; Karakoç et al., 2026). Here we model either no cost or a rather severe cost. This cost is incorporated as a persistently reduced growth rate in the two strategist types, TSS and MSS, which incur the cost constitutively rather than only during switching into dormancy (Karakoç et al., 2026).

The mixed switching strategist (MSS) is described by equivalent equations but with both spontaneous and triggered switching terms included:

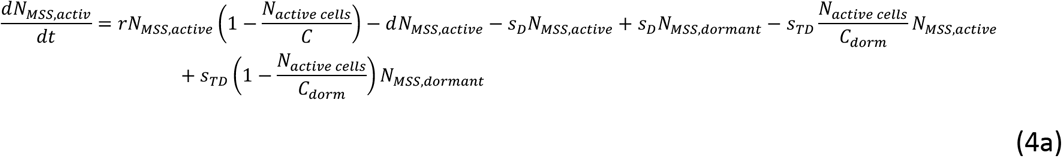

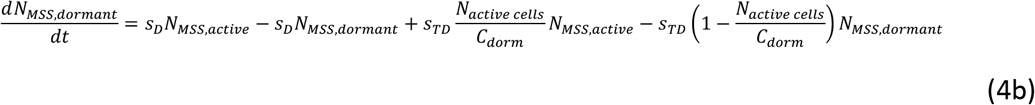

In summary our model has the following features:

First, triggered switching into and out of dormancy is dynamic (to an extent that is tuneable) for both TSS and MSS, and is regulated by competition through niche occupancy, such that the degree to which the niche is filled influences the rate of dormancy induction and reactivation. Second, the model includes a mixed switching strategist (MSS) that switches spontaneously into and out of dormancy while also adjusting its switching rate in response to competition. Third, triggered switching can incur a cost, in the form of a reduced growth rate. Fourth, the model accounts for exploitative competition among the different strategists.

Since our study aims for general insight rather than quantitative predictions, we chose a simple set of parameter values, guided by reported values for soil microbes. The growth rate r was set to a universal value of 0.1/day (Pagel et al., 2016), except in cases where there was a cost for triggered switching, when r was set to 0.05/day, as explained above. The intrinsic death rate was set to 0.0072/day for all active cells while dormant cells are assumed not to die (Joergensen et al., 1990). The spontaneous switching rate s_D_ for the SSS was set to 0.1/day (Pagel et al., 2020) while the triggered switching rate s_TD_ was usually set to 1/day. The carrying capacity, *C*, of the niche was set to 500 to enable simulations over a time range comparable to that of the experimental data from Sorensen et al. Table 1 (Methods section) lists all parameter values.

**Table 1:** Parameters used in the simulations.

| parameter | Simulation of individual strategists | Simulation without costs | Simulation with cost | reference |
|---|---|---|---|---|
| **Capacity C** | 500 | 500 | 500 | Arbitrary |
| **Growth rate r** | 0.1 /day | 0.1 /day | NSS and SSS: 0.1 /day<br>TSS and MSS: r=0.05 /day | Pagel2016 |
| **Spontaneous switching rate** | 0.1 /day | 0.1 /day | 0.1 /day | Pagel2020 |
| **Triggered switching rate** | 1 /day | 1 /day | 1 /day | Arbitrary, see Fig. S4 |
| **Death rate** | 0.0072 /day | 0.0072 /day | 0.0072 /day | Joergensen1990, death rate at 15°C |

A perturbation was modelled by setting *N_i_*_,active_ to zero for all strategists *i*, while leaving *N_i_*_,dormant_ unchanged. A dispersal event was simulated by adding to the existing community, at a specific time point, cells from a predefined community containing active and dormant cells of all strategists (see Methods for details).

The model was simulated numerically to generate predictions for the active and dormant subpopulation dynamics of each strategist (see Methods for details).

## Results

### Different microbial strategists show different growth dynamics and responses to perturbation

To understand the behaviour of our model microbial strategists, we first simulated each strategist in isolation, including a perturbation after 100 weeks that eliminates all active microbes. The non-switching strategist (NSS; Fig. 2A) grows initially exponentially and its population saturates at a size given by C-d. Since all NSS microbes are active, the perturbation eliminates the entire NSS population. For the spontaneous switching strategist (SSS; Fig. 2B), the active subpopulation shows similar dynamics to NSS, i.e. it grows initially exponentially and saturates at C-d. The dormant subpopulation tracks the active subpopulation, with the active and dormant subpopulations being equal in size since the rates of switching into and out of dormancy are equal in our model. In response to the perturbation, the active subpopulation is eliminated, but it is later restored to its former size due to reactivation of dormant microbes and subsequent growth. The SSS is therefore resilient to the perturbation. The triggered switching strategist (TSS; Fig. 2C; here simulated without a sensing cost) shows similar active subpopulation dynamics but different dormant subpopulation dynamics. For TSS, the dormant subpopulation initially remains low, but once the carrying capacity is reached, the dormant subpopulation becomes very large because the rate of switching into dormancy is much higher than the reactivation rate. In response to the perturbation, all active cells are removed, but the active population rapidly recovers due to switching of dormant cells to the active state. The mixed switching strategist (MSS; Fig. 2D) shows behaviour that is intermediate between SSS and TSS; the dormant subpopulation lags the active subpopulation during exponential growth but becomes dominant once the carrying capacity is reached. In response to the perturbation, all active cells are removed, but the active population rapidly recovers through the switching of dormant cells to the active state, like TSS.

**Figure 2:**
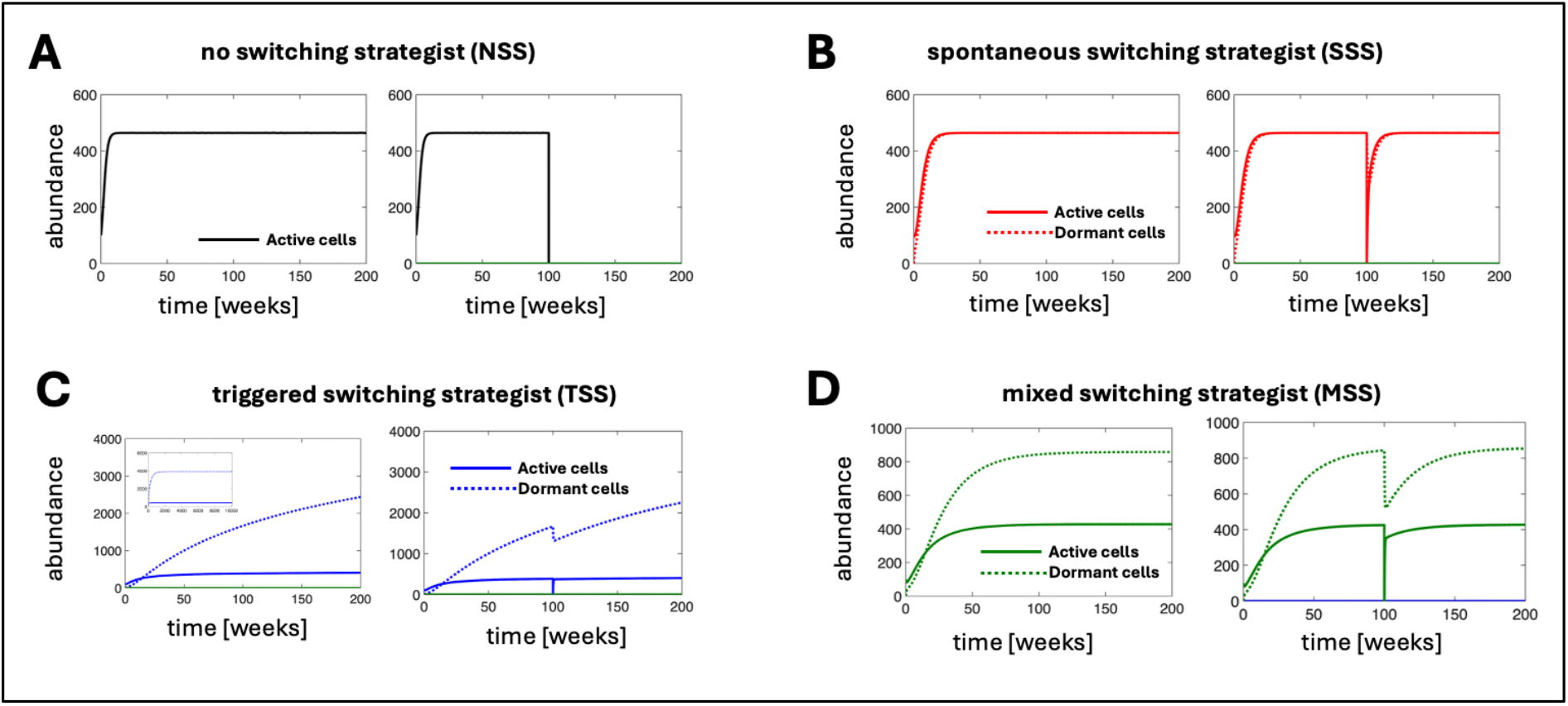
Model dynamics for different strategists, simulated in isolation (left) and with perturbation (right). Each strategist was simulated starting from 100 active microbes (and no dormant microbes) for 200 weeks. The right-hand subfigures show the same simulations as those on the left, but with a perturbation at week 100 that removes all active cells. Note the differences in y-axes ranges across panels. **(A)** For the non-switching strategist (NSS; Eq. (1)), all microbes are active. The NSS grows initially exponentially, and its population saturates at size (C-d). The perturbation eliminates the entire NSS population. **(B)** For the spontaneous switching strategist (SSS; Eqs. (2a,b)), the active subpopulation shows similar dynamics to NSS (panel A). The dormant subpopulation tracks the active subpopulation with the ratio of subpopulation sizes being set by the relative rates of switching into and out of dormancy (here equal). In response to the perturbation, all active cells are removed, but the active population recovers through the switching of dormant cells to the active state. **(C)** For the triggered switching strategist (TSS; Eqs. (3a,b), here simulated without a sensing cost), the active cells again follow the same dynamics as NSS and SSS. The dormant subpopulation lags the active subpopulation but later becomes very large. In response to the perturbation, all active cells are removed, but the active population rapidly recovers through the switching of dormant cells to the active state. **(D)** For the mixed switching strategist (MSS; Eqs. (4a,b); here simulated without a sensing cost), the behaviour is intermediate between SSS and TSS. In response to the perturbation, all active cells are removed, but the active population rapidly recovers through the switching of dormant cells to the active state.

### Switching to dormancy leads to competitive disadvantage in the absence of perturbation

Next, we simulated a mixed community consisting of all 4 microbial strategists, without perturbation or dispersal events. In these simulations, the different strategists compete for the limited niche capacity. Initially we performed these simulations without accounting for a sensing cost. Interestingly, even though the active subpopulations of each strategist showed similar dynamics when simulated in isolation (Fig. 2 A-D), the NSS dominated in competition (Fig. 3A). The SSS strategist had a substantial dormant subpopulation, but even accounting for this, SSS was outcompeted by NSS. For TSS, the active subpopulation was eliminated in competition (even though we did not include a sensing cost). Although there was a substantial dormant TSS subpopulation, the total TSS population was smaller than the total NSS or SSS populations (Fig. S1). Again, the MSS shows behaviour intermediate between SSS and TSS.

**Figure 3:**
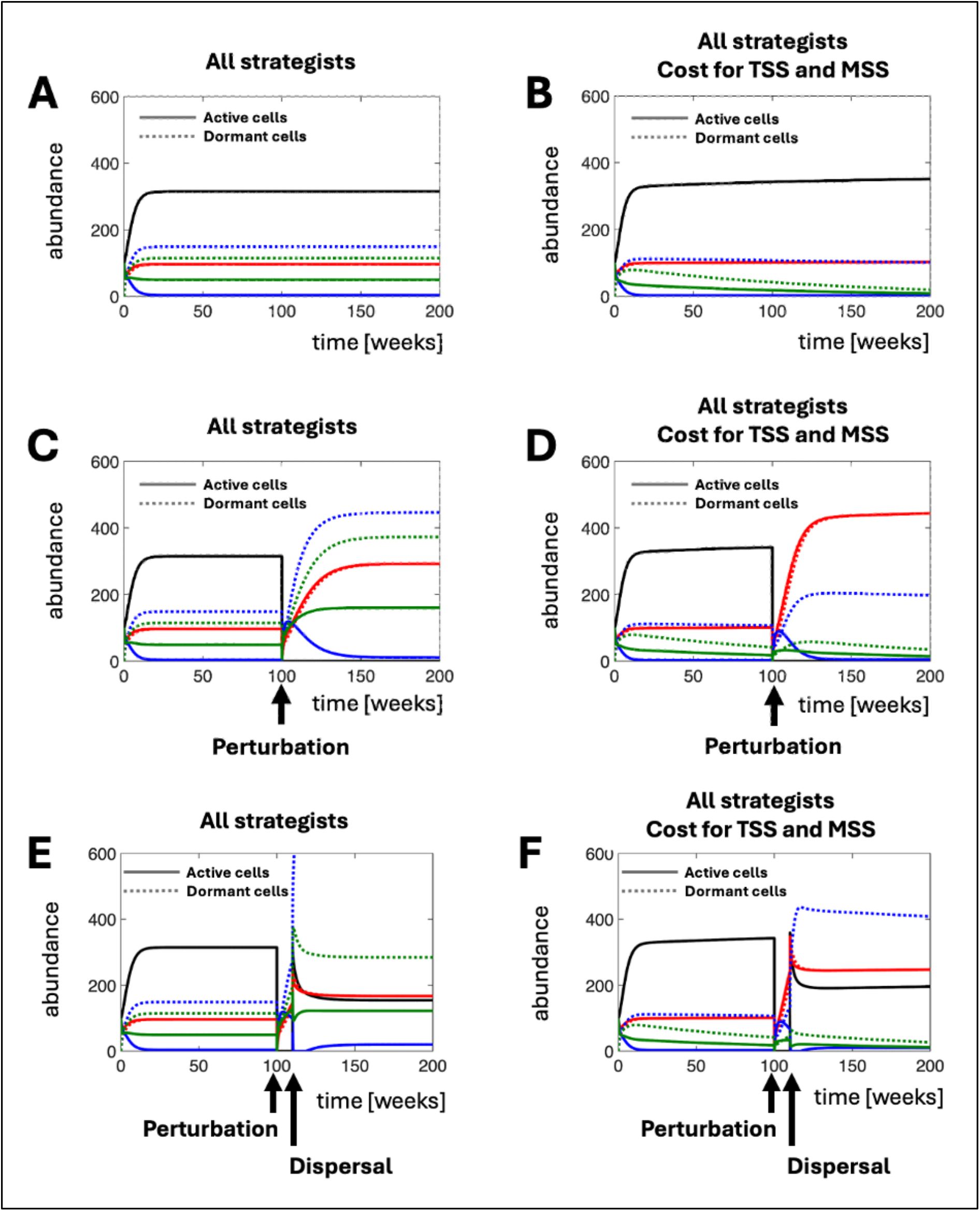
Model dynamics for a mixed community containing all strategists. Simulations were initiated with 100 active microbes of each type (and no dormant microbes) for 200 weeks. The colours show the different strategists: Black: NSS; Red: SSS; Blue: TSS; Green: MSS. Solid lines show active subpopulations; dashed lines indicate dormant subpopulations. **(A)** An unperturbed community, with no sensing cost (i.e. equal growth rate r for all strategists). For SSS the dashed and solid lines coincide. The community becomes dominated by NSS even though all strategists show similar active subpopulation growth in isolation (compare with Fig. 2). **(B)** The same simulation as in (A) but with a sensing cost, i.e. the growth rate r for TSS and MSS is halved. The dominance of NSS is more pronounced with the active subpopulations of TSS and MSS almost vanishing by the end of the simulation. **(C)** A community that is exposed to a perturbation which removes all active microbes. The perturbation is timed such that no strategist has yet been eliminated. NSS are eliminated by the perturbation; the remaining strategists regenerate their active subpopulations by reactivating surviving dormant cells and compete for the niche resources. **(D)** As in (C) but with a sensing cost, i.e. the growth rate r for TSS and MSS is halved. This alters the post-perturbation competitive dynamics. **(E)** A community exposed to both a perturbation that removes all active cells and a dispersal event, in which a mixture of active and dormant cells of each strategist is added from a predefined community. The selected interval of 10 weeks between the perturbation and the dispersal event was guided by the experiment conducted by Sørensen et al. [Sorensen2020]. **(F)** As in (E), but including a sensing cost, i.e. the growth rate (r) of TSS and MSS is reduced by a factor of 2. Under these conditions, NSS and SSS become more dominant, whereas MSS and TSS are nearly eliminated.

These results can be explained by the fact that switching into dormancy imposes a “proliferation cost”; i.e. the transformation of active microbes into dormant ones reduces the effective growth rate of the active subpopulation. Since the different strategists compete for niche capacity, the strategist whose active subpopulation has the highest effective growth rate is expected to dominate. Therefore, a high rate of switching into dormancy effectively reduces the competitiveness of the active subpopulation for niche capacity. This effect is most pronounced for the TSS, which exhibits the highest rate of switching into dormancy when competition for the available niche capacity intensifies.

Next, we repeated our simulations with the inclusion of a sensing cost for triggered switching, represented by decreasing the growth rate of the TSS and MSS strategists, as discussed above. In these simulations, the TSS and MSS were even more disadvantaged (Fig. 3B). By the end of the simulation, the active subpopulations of both TSS and MSS had almost been eliminated. In the case of MSS, the dormant subpopulation also diminished rapidly, consistent with this strategy’s ability to spontaneous switch between active and dormant states. For TSS, the decline in the dormant subpopulation was slower, consistent with its lower rate of reactivation of dormant microbes.

Taken together, these results suggest that switching into dormancy may be disadvantageous within an unperturbed community where competition for resources is strong. This is because, even in the absence of a sensing cost, switching into dormancy reduces the effective proliferation rate of the active microbial subpopulation and hence the ability of the strategist to compete for resources within the niche. This effect is more pronounced if there is a sensing cost associated with switching into dormancy.

### Switching to dormancy provides partial community resilience following perturbation

Dormancy has been hypothesized to function as a bet-hedging mechanism that has a disadvantage under permissive conditions but allows a population to survive harsh conditions (Jones and Lennon, 2010). Linked to this, at the level of a microbial community, previous work has introduced the concept of a “seed bank” of dormant cells that provides resilience, allowing a community to re-establish after a perturbation (Lennon and Jones, 2011). To investigate this in the context of our model, we simulated a mixed community containing all 4 strategists that was exposed to a perturbation event that eliminated all active microbes (Fig. 3 C,D). In these simulations, the NSS strategist was eliminated by the perturbation (as in Fig. 2A) but the other strategists were able to repopulate the niche after the population as their dormant subpopulations re-activated, and the resulting active microbes competed for resources within the niche.

We first performed these simulations without a sensing cost, i.e. with equal growth rate r for all strategists (Fig. 3C). Immediately after the perturbation event, all 3 surviving microbial types (SSS, TSS and MSS) regenerated their active subpopulations due to reactivation of surviving dormant cells. The active subpopulation of TSS grew fastest since this strategist had the largest dormant subpopulation at the time of the perturbation. However the situation changed at later times, when SSS became overall dominant, TSS became mostly dormant and MSS maintained a small active subpopulation and a larger dormant one. Therefore the dynamics at short times was controlled by the relative sizes of the dormant subpopulations (strategists with a larger dormant subpopulation could repopulate the niche faster), but at longer times the competition between different strategists became important, resulting in a community composition that was similar to that before the perturbation, but with the absence of NSS.

When we included a cost for sensing, the picture remained qualitatively similar, but the longer-term dominance of SSS was more pronounced because the sensing cost reduced the competitiveness of TSS and MSS (Fig. 3D). In this case, the relative abundances of SSS, TSS and MSS after the perturbation were significantly different after the perturbation compared with before, probably because the competition between these 3 strategists played out differently in the presence versus absence of NSS.

These results illustrate how dormancy can provide a community with resilience against harsh environmental fluctuations by enabling recovery from dormant cell pools. However, the model suggests that this community-level resilience is partial, and dependent on the different dormancy strategies that are present among community members. Community members that do not have dormant subpopulations cannot reactivate after a mortality-inducing perturbation, and the relative abundances of the surviving community members may also be different following the perturbation, since their interactions play out differently in the absence of the eliminated community members.

### Sensitivity of triggered switching alters the timescale of community response

To assess the effects of gradual triggered switching, compared with a rapid, binary switch of the entire cell pool, we also varied the triggered switching rate s_TD_ (Fig. S4). This did not change the overall outcome: in these simulations, the TSS and MSS strategists still recovered on short timescales post-perturbation due to reactivation of dormant cells, before switching back into dormancy due to increased competition on longer timescales. However, the triggered switching rate s_TD_ did control the timescales over which the TSS and MSS strategists became dormant in response to the competition (Fig. S4). A high triggered switching rate, corresponding to a binary triggered switch into dormancy, caused TSS and MSS to more rapidly enter dormancy, while lower rates allowed TSS and MSS to remain active for longer. This might be relevant in situations where the environment fluctuates in time: here the interplay between environmental timescales and the timescales of dormancy switching may be important (Lastad and Mitarai, 2025).

### Dispersal can push triggered switching strategists into dormancy

Dispersal is an important ecological mechanism that influences microbial community resilience, since immigration can re-introduce taxa that have been eliminated by a perturbation (Sorensen and Shade, 2020, Shade 2024). We modelled dispersal by adding a mixture of active and dormant microbes (Fig. S2) representing all 4 strategists into our simulations at a specific time point. In the model, this corresponds to increasing the values of N_i,active_ and N_i,dormant_ for all strategists i, by a fixed amount at the chosen time. The composition of the added mixture reflected an unperturbed community with the same parameters (see Methods).

We first assessed how well a resident population of each strategist could compete with immigrants that were introduced by a dispersal event. To this end we simulated each strategist individually, then subjected it to a dispersal event where active cells of all types were added (Fig. 4).

**Figure 4:**
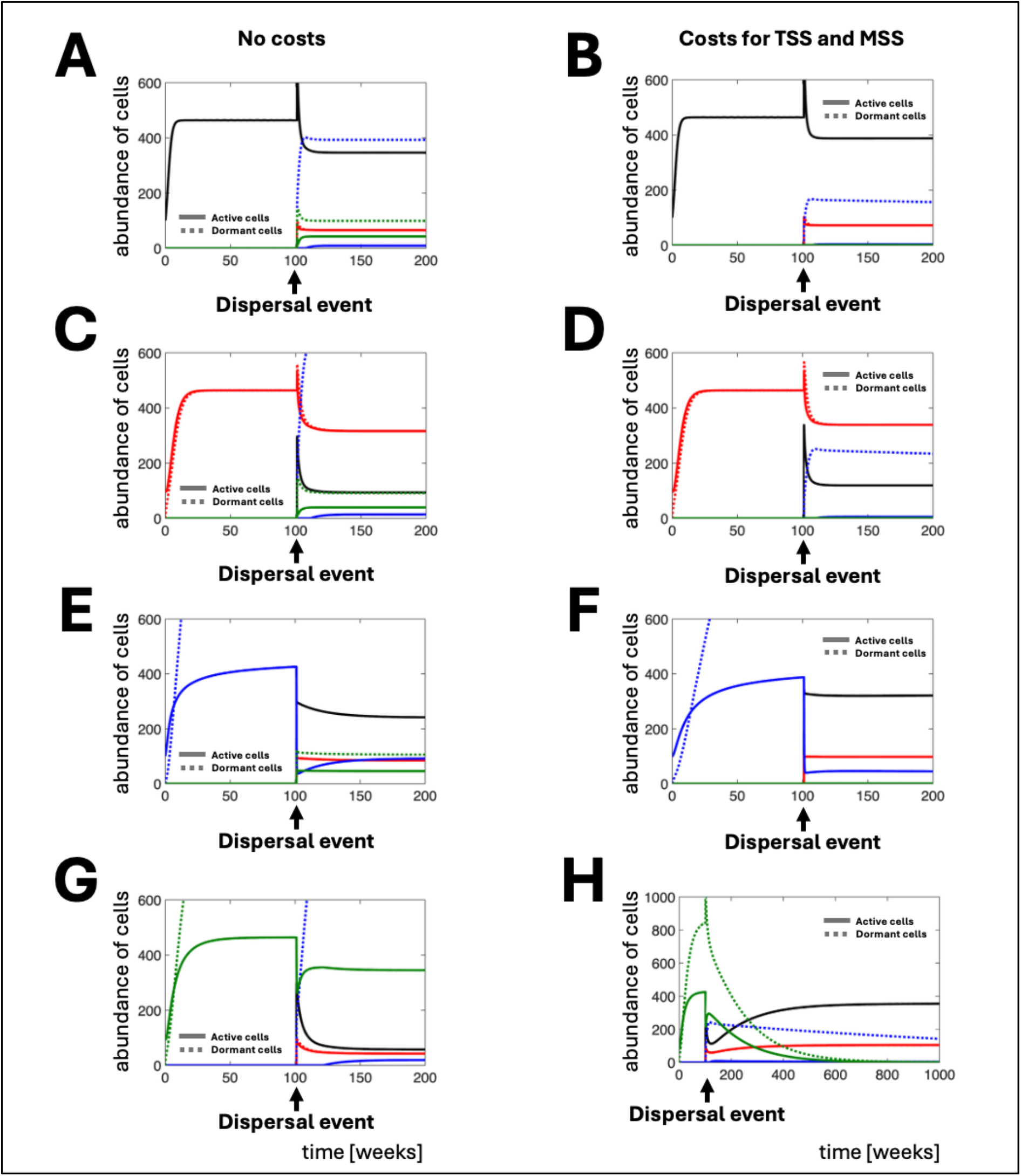
Impact of dispersal on single-strategist resident populations. Single-strategist simulations were initiated with 100 active microbes and were run for 100 weeks before a dispersal event was simulated by adding a mixture of active and dormant microbes of each type from a predefined community. The colours show the different strategists: Black: NSS; Red: SSS; Blue: TSS; Green: MSS. Solid lines show active subpopulations; dashed lines indicate dormant subpopulations. The left panels show simulations without a sensing cost, i.e. with equal growth rate r for all strategists. The right panels show simulations with a sensing cost, i.e. the growth rate r is halved for TSS and MSS. Note differences in y-axes ranges across panels. **(A)** Population of NSS exposed to a dispersal event at week 100, in which a mixture of active and dormant cells of each strategist is added from a predefined community, with no sensing cost (i.e. equal growth rate r for all strategists). The NSS remains the most abundant active strategist in the community following the dispersal event. **(B)** The same simulation as in (A) but with a sensing cost, i.e. the growth rate r for TSS and MSS is halved. The NSS remains the most abundant strategist in the community following the dispersal event and is not surpassed by the dormant TSS. **(C)**Population of SSS exposed to a dispersal event at week 100, in which a mixture of active and dormant cells of each strategist is added from a predefined community, with no sensing cost (i.e. equal growth rate r for all strategists). The SSS remains the most abundant active strategist in the community following the dispersal event. **(D)** The same simulation as in (C), but including a sensing cost, i.e., the growth rate of TSS and MSS is reduced by 50%. Under these conditions, the SSS remains the most abundant strategist and is not surpassed by dormant TSS. **(E)** Population of TSS exposed to a dispersal event at week 100, in which a mixture of active and dormant cells of each strategist is added from a predefined community, with no sensing cost (i.e. equal growth rate r for all strategists). The dispersal event reduces the abundance of active TSS immediately, but during the post-dispersal dynamics, the active TSS pool can recover. **(F)** The same simulation as in (C), but including a sensing cost, i.e., the growth rate of TSS and MSS is reduced by 50%. The active TSS pool is reduced by the dispersal event but does not recover to the same extent as in (E). **(G)** Population of MSS exposed to a dispersal event at week 100, in which a mixture of active and dormant cells of each strategist is added from a predefined community, with no sensing cost (i.e. equal growth rate r for all strategists). The MSS recover rapidly following the dispersal event and become the most abundant active population. **(H)** The same simulation as in (H) but with a sensing cost, i.e. the growth rate r for TSS and MSS is halved. The MSS recover and become the most abundant strategist immediately after the perturbation but become extinct over the long term.

In all cases, the resident strategist was strongly impacted by the dispersal event. For resident NSS or SSS populations the active subpopulation was initially boosted by immigrants but then rapidly declined due to competition for niche resources with immigrants of the other types (Fig. 4A,B,C,D). In the case of SSS, the dormant subpopulation tracked the dynamics of the active subpopulation (Fig. 4B,D). The post-dispersal dynamics of resident TSS or MSS strategists was different (Fig. 4E,F,G,H): here the dispersal event caused an immediate increase in competition which in turn triggered rapid switching into the dormant state. Therefore the dormant resident subpopulation increased after the dispersal event while the active subpopulation decreased.

After the perturbation, community composition was determined by competition between all 4 strategists. This played out differently depending on the identity of the resident type and on the presence or absence of a sensing cost for triggered switching, but as might be expected, competition with immigrants was generally disadvantageous for the resident strategists (Fig. 4).

### Dispersal can restore all strategists following a perturbation

We also investigated the response of mixed communities to dispersal, since previous work shows that dispersal can make an important contribution to the recovery of microbial communities after a perturbation by replacing sensitive taxa whose active subpopulations are eliminated by a perturbation (Sorensen and Shade, 2020), (Shade, 2023).

As above, we simulated a perturbation by removing all active cells from the community and subsequently simulated a dispersal event by adding, at a defined time point, a mixture of active and dormant cells containing representatives of all four strategists and reflecting the composition of an unperturbed community (see Methods). The time interval between the perturbation and the dispersal event was chosen to reflect the experimental design of (Sorensen and Shade, 2020). We repeated these simulations with and without a cost for triggered switching.

In both scenarios, with and without a cost for triggered switching, the NSS population, which was eliminated by the perturbation, was able to recover following dispersal (Fig. 3 E+F). In general the dispersal event increased niche occupancy, leading to an immediate increase in competition among strategists. Consequently, the competition-sensitive TSS and MSS rapidly reduced their active populations by switching into dormancy. Therefore our simulations are consistent with previous suggestions that dispersal contributes to resilience after a perturbation by replacing eliminated community members (Sorensen and Shade, 2020), (Shade, 2023), but they also demonstrate a key role for exploitative competition (i.e. competition for niche resources) between immigrant and resident community members, that may play out differently for different dormancy strategists.

### Model-derived hypotheses are partially consistent with a dynamic soil community dataset

Taken together, our model results suggest four hypotheses for how microbial community activity dynamics may respond to perturbation and dispersal events: (i) active and dormant subpopulations are expected to coexist stably, due to reversible switching between active and dormant states; (ii) If dormant microbes are tolerant to a perturbation, dormant subpopulations can reactivate after a perturbation, providing partial community resilience, even though dormancy can be a competitive disadvantage in the absence of perturbation; (iii) a dispersal event may push some microbial types into dormancy due to increased competition with immigrant microbes, (iv) a dispersal event can restore eliminated community members after a perturbation, contributing to community resilience.

These hypotheses were derived from a highly idealised model. To assess whether their validity in natural, complex microbial communities, we reanalysed the dataset of Sorensen et al., who tracked community-level activity dynamics in heat-perturbed and dispersed soil mesocosms (Fig. 5) (Sorensen and Shade, 2020). In the experiments of Sorensen et al., subsets of mesocosms were either maintained at 14°C for 45 weeks (as controls), subjected to a 2-week heat perturbation at 60°C at weeks 4-6 and allowed to recover at 14°C for a total of 45 weeks, or subjected to the heat perturbation followed by a dispersal event in which a small sample from an unperturbed community was introduced (see Methods for details). The active and dormant microbial community composition was tracked over time by 16S ribosomal RNA and RNA gene amplicon sequencing.

**Figure 5.**
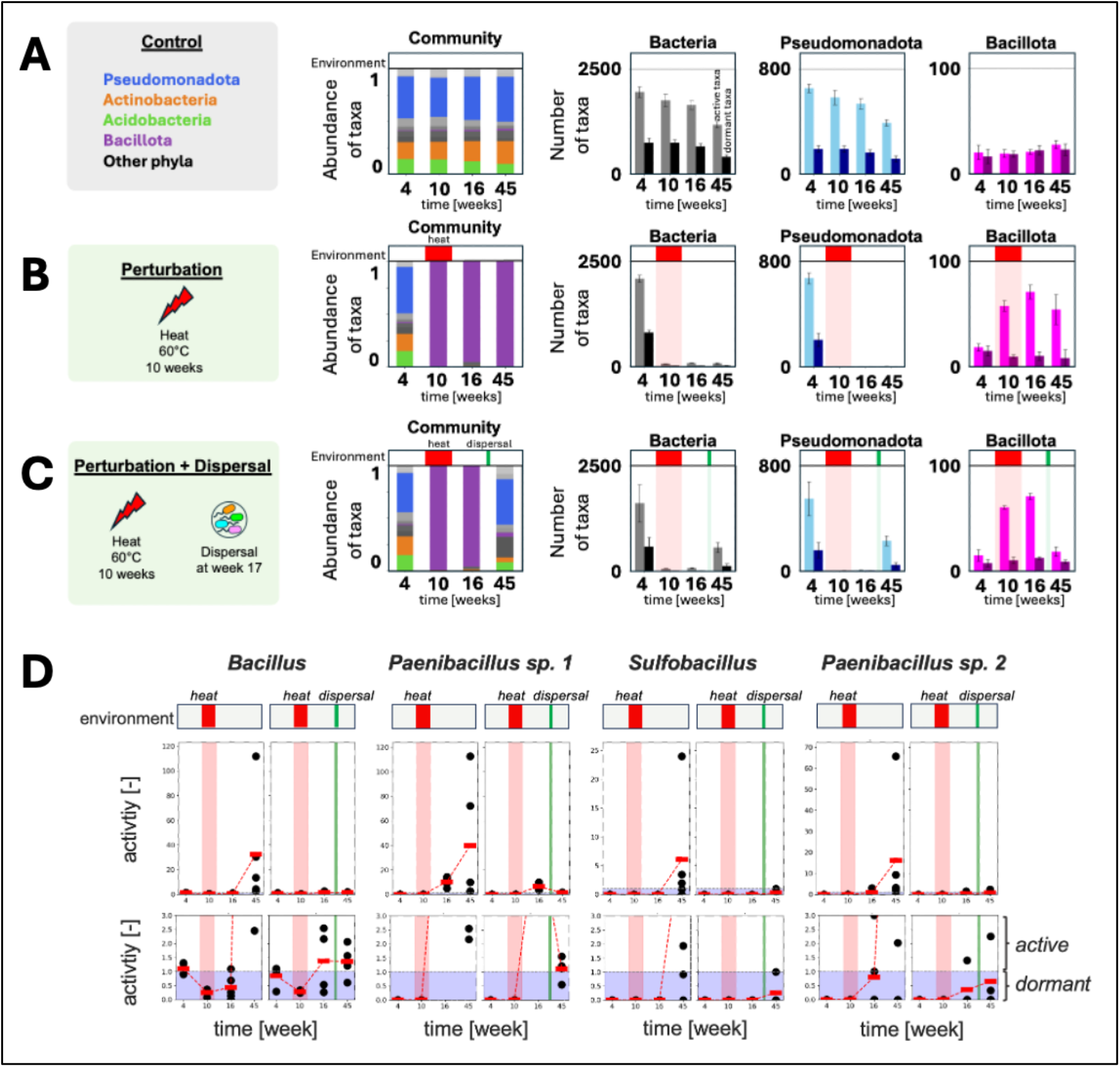
Summary of reanalysed experiment from (Sorensen and Shade, 2020). (A+B+C) Microbial community compositions in three soil mesocosms: (A) an unperturbed control, (B) a perturbed mesocosm, and (C) a perturbed mesocosm with dispersal. **(A)** A control mesocosm. Left: Relative abundance of taxa corresponding to different phyla over time, as determined by 16S rRNA gene analysis. Right: the numbers of taxa that were classified as active (lighter bars) or dormant (darker bars), at different times during the experiment, plotted for (i) all bacteria, (ii) Pseudomonadota and (iii) Bacillota. Classification into active and dormant taxa was based on rRNA vs rRNA gene 16S amplicon abundances, see Results. **(B)** Equivalent data for a heat-perturbed mesocosm without dispersal. (C) Equivalent data for a heat-perturbed mesocosm with dispersal. In (B) and (C) the red and green shading indicate the timing of the heat perturbation and dispersal event, respectively. **(D) Experimentally measured activity profiles of different taxa of a soil mesocosm kept for 45 weeks under lab conditions.** Bottom row is a zoom in to show activity changes at low activity values. Some mesocosms were treated with a heat perturbation (red area) and additionally with a dispersal event (green line). The left subfigures received only heat perturbation. The right subfigures received a heat perturbation and afterwards, a dispersal event. Activity of different taxa was obtained by measuring the 16S RNA/16S rRNA gene amplicons at 4, 10, 16, and 45 weeks. Activity value between 0 and 1 indicates dormant state, value higher than 1 indicate active state. Black dots are single measurements; red bar indicates mean value. Data from (Sorensen and Shade, 2020).

Starting from Sorensen et al.’s dataset of 16S ribosomal RNA and rRNA gene counts, we re-analysed the observed community dynamics in the light of our model-derived hypotheses.

### Control mesocosms maintain stable microbiomes with significant dormancy

Motivated by the model hypothesis that reversible active-dormancy switching leads to stable dormant pools, we assessed whether stable dormant subpopulations were maintained over long times in control mesocosms that were not subject to perturbation or dispersal. 16S rRNA amplicon analysis showed that the control mesocosms contained diverse bacterial communities dominated by Pseudomonadota (formerly Proteobacteria) with significant abundances of Actinomycetota (formerly Actinobacteria), Acidobacteriota (formerly Acidobacteria) and Bacillota (formerly Firmicutes); community composition (at the phylum level) remained stable through the entire experiment (Fig. 5A). To analyze the dormant sub-community, individual taxa (OTUs) were classified as being either active or dormant based on the abundances of their rRNA and rRNA gene amplicons (see Methods). Based on this analysis, all the major phyla contained both active and dormant taxa, in different proportions (Fig. 5 A). For example, most Proteobacterial taxa were classified as active, while about half of the detected Bacillota taxa were classified as dormant, reflecting also their different natural histories, with major Bacillota lineages being prominent spore-formers. For all the abundant phyla, the proportion of dormant taxa appeared to be stable over long times (Fig. 5A) suggesting that the dormant pool in these mesocosms was indeed stable.

### Thermotolerant taxa survive heat perturbation but dormancy does not provide protection

In our model, we assumed that dormant microbes are protected from a perturbation, consistent with the fact that some dormant states, such as spores, are known to provide tolerance to environmental perturbations (Setlow, 2014b). We did not find evidence for this scenario in the Sorensen et al. dataset. As previously shown, in this experiment heat perturbation caused a drastic shift in community composition (Sorensen and Shade, 2020). During and after the perturbation the Pseudomonadota along with the Actinomycetota and Acidobacteriota, essentially disappear and the mesocosms become almost completely dominated by Bacillota, that were previously at relatively low abundance (Fig. 5B). Interestingly, both active and dormant taxa were eliminated by the perturbation, both for the bacterial community, and for the Proteobacteria in particular, suggesting that dormancy did not, in general, provide protection. We note, however, that some taxa may have survived the perturbation in a dormant state and not been detected by the RNA/DNA-based method, if their abundance was below the detection limit.

Among the thermotolerant Bacillota taxa, both active and dormant taxa were able to survive the perturbation. The number of active Bacillota taxa increased during the perturbation and remained high after the perturbation (Fig 5B). However, this increase could not be ascribed to a net shift from dormancy to activity, since the number of dormant Bacillota taxa remained stable through the experiment (Fig. 5B). Rather, previously undetectable Bacillota taxa may have increased in abundance and become detectable during the perturbation. This data is consistent with a picture in which Bacillota taxa profit from the heat perturbation by being able to survive when others are eliminated, and by expanding to occupy the vacated niche – dynamics which mirror phenomena that we observed in our model (e.g. Fig. 3C,D). However, since we did not find evidence that dormant taxa preferentially survived the heat perturbation (rather, the survivors were both active and dormant thermotolerant Bacillota taxa), the model assumption that dormancy provides protection may not be valid for this dataset. This highlights the importance of considering the specific characteristics of the perturbation, particularly its intensity and the extent to which it imposes strong selective pressure, when evaluating the role of dormancy in community resilience.

### Dispersal increases competition and can push some taxa into dormancy

In our model, introduction of immigrant microbes via a dispersal event increased competition for resources, which triggered a switch to dormancy for some strategists. Interestingly, Sorensen et al.’s dataset may be consistent with competition following the dispersal event. Even though the amount of dispersed material was very small (less than 0.1 % dispersed volume to received volume), dispersal had a strong effect on community composition (Fig. 5C). In particular, the Bacillota that had dominated after the perturbation were suppressed, suggesting possible increased competition for the niche that they had occupied immediately after the perturbation (Fig. 5C, right; compare with Fig. 5B).

Interestingly, the number of detected active Bacillota taxa decreased strongly after the dispersal event, while the number of detected inactive Bacillota taxa remained relatively unchanged (Fig. 5C, right; compare with Fig. 5B). To understand whether individual active taxa might have switched into dormancy in response to the dispersal event, as opposed to simply becoming undetectable, we tracked the ratio of 16S rRNA to 16S rRNA gene reads (as a measure of activity level) for individual Bacillota taxa through the experiment (Fig 5D). We identified several Bacillota taxa, including representatives of *Bacillus*, *Sulfobacillus* and *Paenibacillus*, that responded to dispersal by decreasing the level of their activity (Fig. 5D). Therefore, this analysis supports the model-derived hypothesis that dispersal can lead to increased competition and may push some community members towards inactivity.

### Dispersal contributes to resilience by restoring sensitive members of the soil community

In our model, a dispersal event can restore eliminated community members after a perturbation, contributing to community resilience. Consistent with this concept, in Sorensen et al.’s experiments a dispersal event was followed by a strong increase in microbial diversity, characterised by an increase in the overall number of detected bacterial taxa (Fig. 5C). Those phyla that had been largely eliminated by the perturbation (in particular, Pseudomonadota, Actinomycetota and Acidobacteriota) were able to partially recolonize the mesocosms after the dispersal event, reflected in an increase in their number of detected taxa (e.g. Fig. 5C for Pseudomonadota). As previously noted (Sorensen and Shade, 2020), the final community composition of the dispersal-treated mesocosms was much closer to the pre-perturbation composition than was that of the perturbed mesocosms without dispersal (Fig. 5C), suggesting that dispersal contributed to resilience by restoring some thermosensitive community members that could successfully compete for niche resources with members of the post-perturbation community. However, also consistent with our model, community resilience was only partial (community composition was significantly different pre- and post-perturbation even with dispersal (Sorensen and Shade, 2020)).

## DISCUSSION

### Exploitative competition among dormancy switching strategists can shape microbial community response to perturbation and dispersal

Dormancy is widespread among plants, animals and in microbial communities such as the soil microbiome, yet its ecological role remains a topic of discussion. Previous theoretical microbial ecology studies have focused on dormancy as a possible bet-hedging mechanism in the face of perturbations (linked at the community level to the seed bank hypothesis), or as a mechanism for avoidance of competition (Gardner et al., 2007). Our work suggests that the ecological implications of dormancy at the community level may involve a complex interplay between the individual-cell protection from perturbation that may be provided by some forms of dormancy, and competition for niche resources among different dormancy switching strategists within the community. Different dormancy switching strategies could result in differently sized active cell pools (see, e.g. Figure 2) and provide different levels of protection from perturbation. Different strategies also have different competitive implications: for example, in our model, strategists possessing triggered switching (TSS and MSS) were predominantly maintained in a dormant state when embedded in a microbial community, but not when simulated in isolation (Figure 3, compare to Figure 2). Our model suggests that different dormancy strategies can interact in multiple ways, including through exploitative competition, potential competition-triggered switching and preferential survival of perturbations. These interactions can shape community dynamics across multiple timescales, from immediate responses to perturbations to long-term shifts in the relative sizes of active and dormant populations, ultimately influencing community composition and resilience.

### Transient niche occupation after a perturbation is followed by competition that is increased by dispersal

Our model suggests that the response of a microbial community to a severe perturbation may play out differently on short versus longer timescales. In our model, a perturbation was assumed to remove all active cells from a microbial community. On short timescales, surviving dormant cells that were present before the perturbation were able to reactivate and rapidly repopulate the vacant niche; during this time the carrying capacity of the niche was not fully occupied, resulting in reduced competition among the active subpopulations of the survivors (Figure 3). However, on longer timescales, as the niche became occupied, competition intensified, causing strategists with triggered switching mechanisms to return to the dormant state. A dispersal event increased this competition by re-introducing sensitive strategists that were removed by the perturbation. Even though not all aspects of our model were confirmed in our analysis of the experimental dataset of Sorensen et al (2020) (specifically, we did not see evidence for increased survival of the perturbation among dormant taxa), the rapid population of the niche by survivors of the perturbation, followed by their apparent competition-driven replacement following the dispersal event, were features of the experimental dataset (Figure 5). Indeed, the potential of a dispersal event to increase competition, displacing less competitive microorganisms that were able to repopulate the niche after the perturbation, is an important insight of our model. This effect may be relevant in the context of microbial inoculation strategies aimed at improving soil functions, where introduced microorganisms may alter the activity states and persistence of resident microbial communities (King and Bell, 2022).

### Costs of dormancy switching and their ecological implications

Previous work has assumed that a triggered dormancy strategy is associated with intrinsic costs compared with a stochastic switching strategy; these costs are thought to arise from the energetic requirements for building and operating a sensing mechanism (Govern and Ten

Wolde, 2014; Karakoç et al., 2026; Kussell et al., 2005). In these previous models, the costs of triggered switching are balanced by the benefits of early activation or rapid entry into dormancy to survive harsh environmental conditions (Govern and Ten Wolde, 2014). To account for this, we compared the predictions of our model in the presence and absence of a cost of triggered switching, in the form of a severely reduced growth rate for the TS and MS strategists (Bruggeman et al., 2020). In the absence of costs for triggered switching, different switching strategists were able to coexist within the same community (Figure 3). However, when costs were included, the MSS, with both spontaneous and triggered switching mechanisms, could not be maintained in a stable microbial community. We note that the magnitude of the cost for triggered switching was here chosen to be very high (in the form of a halving of the growth rate); further experimental work would be needed to determine realistic costs for triggered switching for different species and further simulations would be required to determine the effect of the cost magnitude parameter within a microbial community context.

Our results may imply that simultaneous operation of both stochastic and costly triggered switching mechanisms may not be a stable strategy, especially at lower rates of triggered switching. A mixed switching strategy has previously been discussed in the context of the ‘scout’ hypothesis (Buerger et al., 2012; Epstein, 2009), that suggests that some cells within a population may act spontaneously (the scouts), whereas others switch upon receiving a trigger from the scouts - however our model did not encompass the triggering of activation of one subpopulation by another. Further work accounting for individual cell heterogeneity (within each strategy group) would be needed to understand the implications of the scout hypothesis for population dynamics within the framework of our model. It is also possible that temporal partitioning of MSS, whereby the entire population switches between spontaneous and triggered dormancy depending on prevailing conditions, might stabilize this strategy. Such plasticity could allow the MSS to combine the benefits of spontaneous and triggered mechanisms across fluctuating environments (Balaban et al., 2004).

A further insight arising from our model relates to the growth opportunity costs of dormancy switching. Even when there is no explicit cost associated with the switching mechanism, switching into dormancy reduces the effective growth rate because active microbes are transformed into dormant cells, that do not proliferate. This phenomenon is closely linked with the previously discussed bet-hedging trade-off between maintaining a dormant state and maximizing growth: while dormancy may help a population survive a perturbation, it comes at the expense of a reduced effective growth rate (Kussell and Leibler, 2005).

### Different strategies lead to different dormant cell pool sizes

Our model also highlighted a further ecological implication of stochastic versus triggered switching strategies: these strategies produce different characteristics of the dormant cell pool. Stochastic switching strategists (SSS) maintain a dormant pool that tracks the active cell pool (at least for reversible stochastic switching, as modelled here). Thus, a spontaneous switching strategy may allow for maintenance of an active population capable of withstanding competition as well as a dormant population that can serve as a reservoir in the event of active-cell extinction. However the SSS pays an opportunity cost in the form of reduced total proliferation in the absence of a perturbation: the active cell pool for SSS is smaller than that for the NSS. In contrast, in our model the TSS were found to remain predominantly in the dormant state when in a mixed community, because we assumed that competition for resources triggered the switch to dormancy. This resulted in a large dormant pool for TSS, allowing them to very rapidly repopulate the niche following the perturbation; however, they were weak competitors (Figure 3D). This picture may be specific to the resource-competition trigger that we assumed here; TSS that respond to other triggers might be able to maintain a larger active pool within a mixed community and hence be stronger competitors. In general our model makes clear that the presence of triggered switchers, and the environmental parameters that they respond to, changes the total dormant pool size and its environmental responsiveness. Hence, dormant pool size reflects both environmental conditions (present and past) and the relative abundance of different strategists within the community.

### Limitations of the model

Our model is idealised both in its choice of mechanisms (e.g. the resource-competition driven trigger for TSS, and the elimination of active but not dormant cells by the perturbation), and in its parameter values. It also does not capture the behaviour of individual cells within each strategy group; this may be especially relevant for triggered switching, where the gradual but homogeneous response of the entire population that we modelled here may have different ecological implications compared to a scenario where individual cells switch at different times in response to the trigger. In general, our model aims to provide conceptual insights into possible ecological implications of dormancy switching strategies within a mixed community subject to perturbation and dispersal, rather than to provide either quantitative or qualitative predictions. Despite its limitations, our model highlights how ecological outcomes may depend on a complex interplay between dormancy switching, resource competition, perturbation and dispersal.

### Soil mesocosm dataset generally supports model insights

Our re-analysis of the soil mesocosm experiment of Sorensen and Shade (2020) was generally supportive of our model’s main insight that community response to perturbation and dispersal depends on an interplay between dormancy and exploitative competition. However, there was a clear discrepancy between the model and the data analysis in that we did not find evidence that dormancy provided general protection from the heat perturbation. Rather, thermotolerant taxa appeared to survive the perturbation independent of their dormancy status. Our analysis was, however, limited in that taxa were scored as being either entirely active or entirely dormant. We did not take account of quantitative activity levels, and some taxa may have become undetectable in our analysis even if they were still present in the experiment. Another important limitation was that our data consisted of relative rather than absolute abundances of community members. Therefore, we cannot assess whether the replacement of thermotolerant taxa by immigrant thermosensitive taxa after the dispersal event (Figure 6) occurred in conditions of strong or weaker resource competition.

### Outlook

Mathematical models have the potential to disentangle the complex interplay between dormancy, competition, perturbation and dispersal, and hence to predict and understand the dynamics of microbiomes in changing environments. To do so, however, more information is required on how individual taxa switch to and from dormancy, what triggers are relevant, what the costs are, which dormant states provide protection from which perturbations, the extent of individual cell heterogeneity in dormancy behaviour within taxa, and the extent to which different activity phenotypes might be more or less likely to disperse. Advances in methods such as microscopic identification of individual dormant cells within communities may soon provide such information and may be combined with individual-based modelling (Bäcker et al., 2026).

We evaluated our model-derived insights using an experimental dataset from a soil microbial community. To determine how broadly these insights apply, future studies could investigate taxon-level changes in activity levels in response to perturbations and dispersal in natural environments such as aquatic systems, sediments, and host-associated microbiomes, as well as in managed microbial systems, including wastewater treatment, bioremediation, agriculture, and industrial bioprocesses. These systems differ in community composition, environmental conditions, dispersal rates, and the type and intensity of perturbations, all of which may influence dormancy and reactivation. Further cross-system testing would help establish whether the predicted interplay between dormancy, exploitative competition, and response to perturbation and dispersal represents a general principle of microbial community resilience. Such knowledge could inform management strategies that use microbial seed banks or targeted dispersal to maintain or restore important ecosystem and biotechnological functions.

## CONCLUSION

In this study, we used a niche-based mathematical model to show how dormancy-switching mechanisms and exploitative competition within a microbial community can shape the community’s response to perturbations and dispersal. Our model, comprising four microbial strategists that differ in their use of spontaneous and triggered switching into and out of dormancy, allowed us to investigate the dynamics of active and dormant cell pools under stable conditions, perturbation, and dispersal. We also compared models with and without triggered switching costs and with gradual vs binary triggered switching.

Our model simulations, combined with a re-analysis of a previous soil mesocosm dataset, support previous findings that perturbation-tolerant dormant cells can reactivate after disturbance and thereby contribute to population recovery, and that dispersal can restore community members that have been eliminated by perturbation and thus promote community resilience. Our results also contribute a new perspective, that dispersal can increase competition for niche resources and hence drive some resident community members into dormancy. Taken together, these findings show how an interplay between exploitative competition and the different dormancy strategies that are present within a microbial community can shape the response of the community to perturbations and dispersal.

## METHODS

### The model and its implementation

The model simulates the population sizes *N_i_*_,state_, for microbial strategist *i* ∈ {1, 2, 3, 4} in either the active or dormant metabolic state. Active cells are assumed to grow at a growth rate *r* and die at a death rate *d* (Fig. 1). Dormant cells do not grow or die.

We adopted a Lotka–Volterra-style competition model (Altermatt et al., 2011; Gonze et al., 2018), in which each active or dormant subpopulation is represented by an ordinary differential equation. The equations are presented above. All strategists were assumed to be initially in the active state, i.e., initial values of N_i,active_ were set to 100 and N_i,dormant_ were set to 0. The system of ODEs was implemented in MATLAB and solved numerically using a standard ODE solver (ode45). The model tracks the seven possible state variables corresponding to the active and dormant fractions of each type.

To account for the possible extinction of a subpopulation within our simulations, any values for *N_i_*_,state_ falling below 1 (assuming cell numbers) were set to zero during the simulation. This threshold was applied to both active and dormant subpopulations.

The parameter values used in our model were chosen to loosely correspond to soil communities. Parameters were derived from available data if possible and based on logical consideration elsewhere. For simplicity, parameter values were assumed to be the same across different strategists where possible. Table 1 lists the parameters used.

A dispersal event was modelled by adding a predefined community composed of active and dormant cells (Fig. S2) at a specified time point. The predefined community consists of cells from a simulated community modelled with the same parameter values. The predefined community was initiated with 100 active cells of each strategist and simulated for 1000 weeks. After 1000 weeks, the community without costs for TSS and MSS had reached a steady state, but the community with costs for TSS and MSS had not yet reached a steady state: the MSS had already been eliminated, whereas the TSS persisted. Hence this represents an intermediate state between the short-term situation, in which all strategists are present, and the long-term outcome, in which both MSS and TSS are eliminated.

When the predefined community was added to a similar community, we observe transient dynamics of active cells, which eventually return towards their initial abundance, while the dormant pool becomes enriched with dormant cells, particularly TSS (Fig. S3). Hence, combining two similar communities has an effect on the dormant pool size.

Specifically, to model dispersal in simulations without costs for triggered switching, the added number of cells was: NSS (only active: 315), SSS (active: 96; dormant: 96), TSS (active: 3; dormant: 149), MSS (active: 49, dormant: 115). For the simulations with costs for triggered switching the number of added cells was: NSS (only active: 359), SSS (active: 104; dormant: 104), TSS (active: 2; dormant: 65), MSS (active: 0, dormant: 0).

### Re-analysis of experimental data

In the experiments performed by Sorensen et al., 15 x 300mL mesocosms of soil from Centralia, PA, USA, were maintained for 45 weeks under different regimes of heat perturbation and dispersal (Sorensen and Shade, 2020). All mesocosms were initially maintained at 14°C for four weeks. Nine of the mesocosms were then subjected to heat disturbance: the temperature was gradually increased to 60°C over two weeks, maintained at 60°C for 8 weeks and decreased again over two weeks to 14°C. To mimic a dispersal event, four of these nine mesocosms were inoculated with a small sample (less than 0.1 % dispersed volume to received volume) of an unperturbed control mesocosm at week 17. The remaining six mesocosms were maintained at 14°C for the entire experiment, as a control. At specific time points, mesocosms were sampled (by removal of 15 g soil) and the active and dormant microbial community composition was analyzed by co-extraction of DNA and RNA followed by amplification and Illumina sequencing of 16S rRNA and rRNA gene amplicons (Sorensen and Shade, 2020).

The values in the data set used to re-analyze the experiment, are the numbers of reads. The rRNA gene represent the relative abundance of an individual operational taxonomic unit (OTU), and the cDNA (RNA) values represent the detected ribosome numbers per OTU. We define active community members with 16S rRNA:rRNA gene ratio > = 1, meaning that there were many more counts of ribosomes per cell than rRNA genes detected. Dormant OTUs were defined with a 16S rRNA:rRNA gene ratio <1 and >0, as well as for rRNA gene value equals 0 and rRNA value >0. Each taxon was classified originally using SINTAX and with the Silva database (Sorensen and Shade, 2020), and here we have manually updated the taxonomic nomenclature of previously identified taxa to reflect recent updates (Oren et al., 2023). The rRNA gene values were used to calculate the relative abundance of each OTU within the community.

## Acknowledgments

AU and RJA were supported by the the Deutsche Forschungsgemeinschaft (DFG) via the Excellence Cluster Balance of the Microverse (EXC 2051 - Project-ID 390713860). AS acknowledges support from the European Union (ERC, MicroRescue, 101087042). Views and opinions expressed are however those of the author(s) only and do not necessarily reflect those of the European Union or the European Research Council. Neither the European Union nor the granting authority can be held responsible for them.

## Supplements

**Figure S1:**
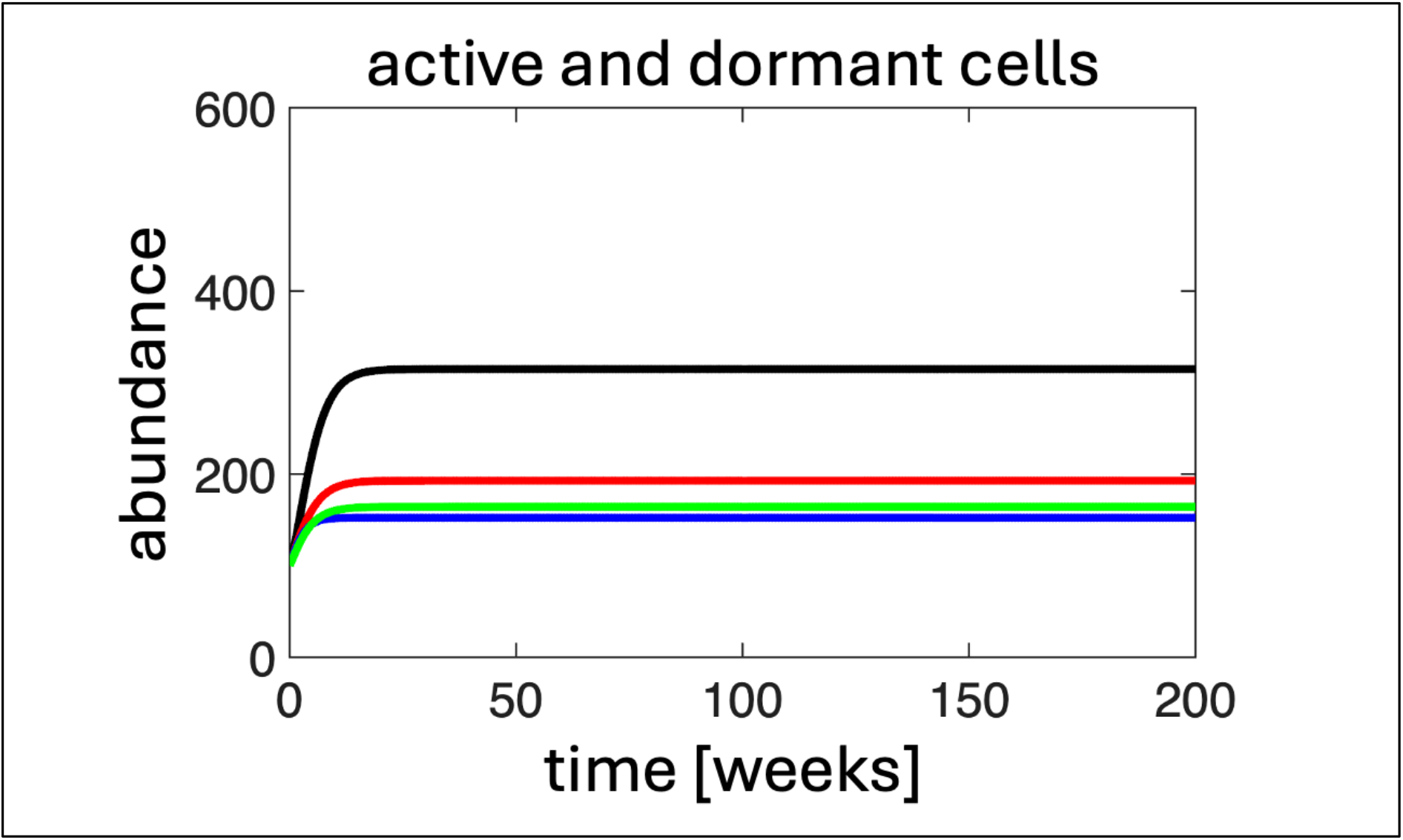
Simulation was initiated with 100 active microbes of each type (and no dormant microbes) and simulated for 200 weeks. TSS and MSS without costs, i.e. halved growth rate. The colours show the different strategists: Black: NSS; Red: SSS; Blue: TSS; Green: MSS. Solid lines show total population (actives+dormants) for each strategist. x-axis: time in weeks; y-axis: abundance.

**Figure S2:**
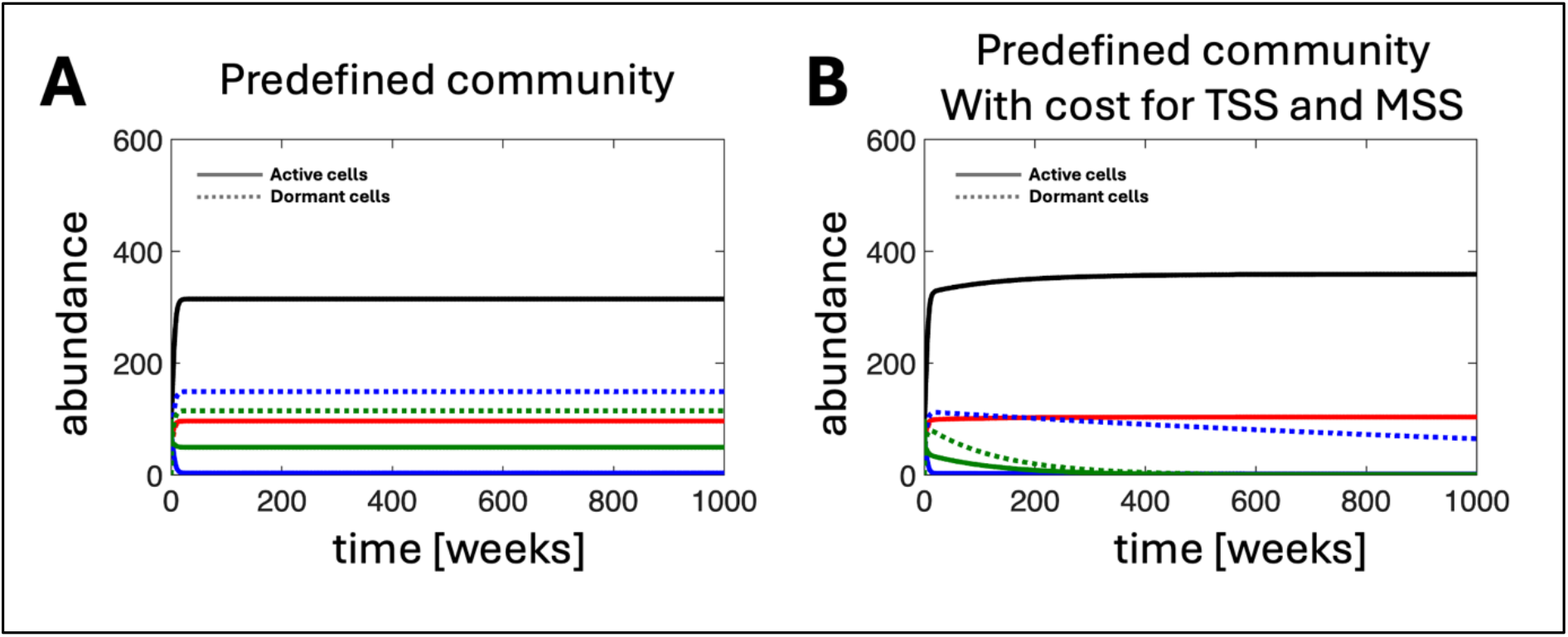
Generation of a community used for dispersal. Simulation was initiated with 100 active microbes of each type (and no dormant microbes) and simulated for 1000 weeks such that the MSS was not yet eliminated. The colours show the different strategists: Black: NSS; Red: SSS; Blue: TSS; Green: MSS. Solid lines show active subpopulations; dashed lines indicate dormant subpopulations. x-axis: time in weeks; y-axis: abundance.

**Figure S3:**
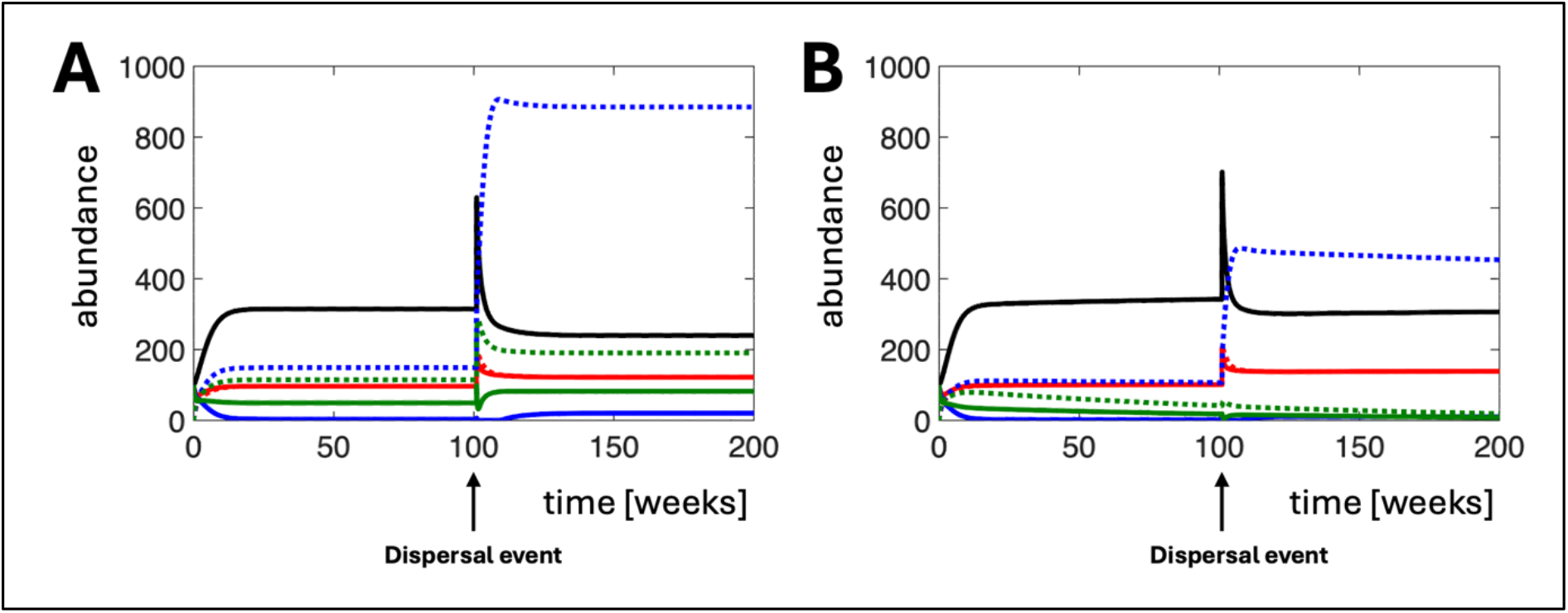
Model dynamics for a mixed community containing all strategists and exposed to a dispersal event. Simulations were initiated with 100 active microbes of each type (and no dormant microbes) and simulated for 200 weeks. After 100 weeks a dispersal event introduces active and dormant cells from a predefined community. The colours show the different strategists: Black: NSS; Red: SSS; Blue: TSS; Green: MSS. Solid lines show active subpopulations; dashed lines indicate dormant subpopulations. **(A)** An unperturbed community, with no sensing cost (i.e. equal growth rate r for all strategists). **(B)** The same simulation as in (A) but with a sensing cost, i.e. the growth rate r for TSS and MSS is halved. x-axis: time in weeks; y-axis: abundance.

**Figure S4:**
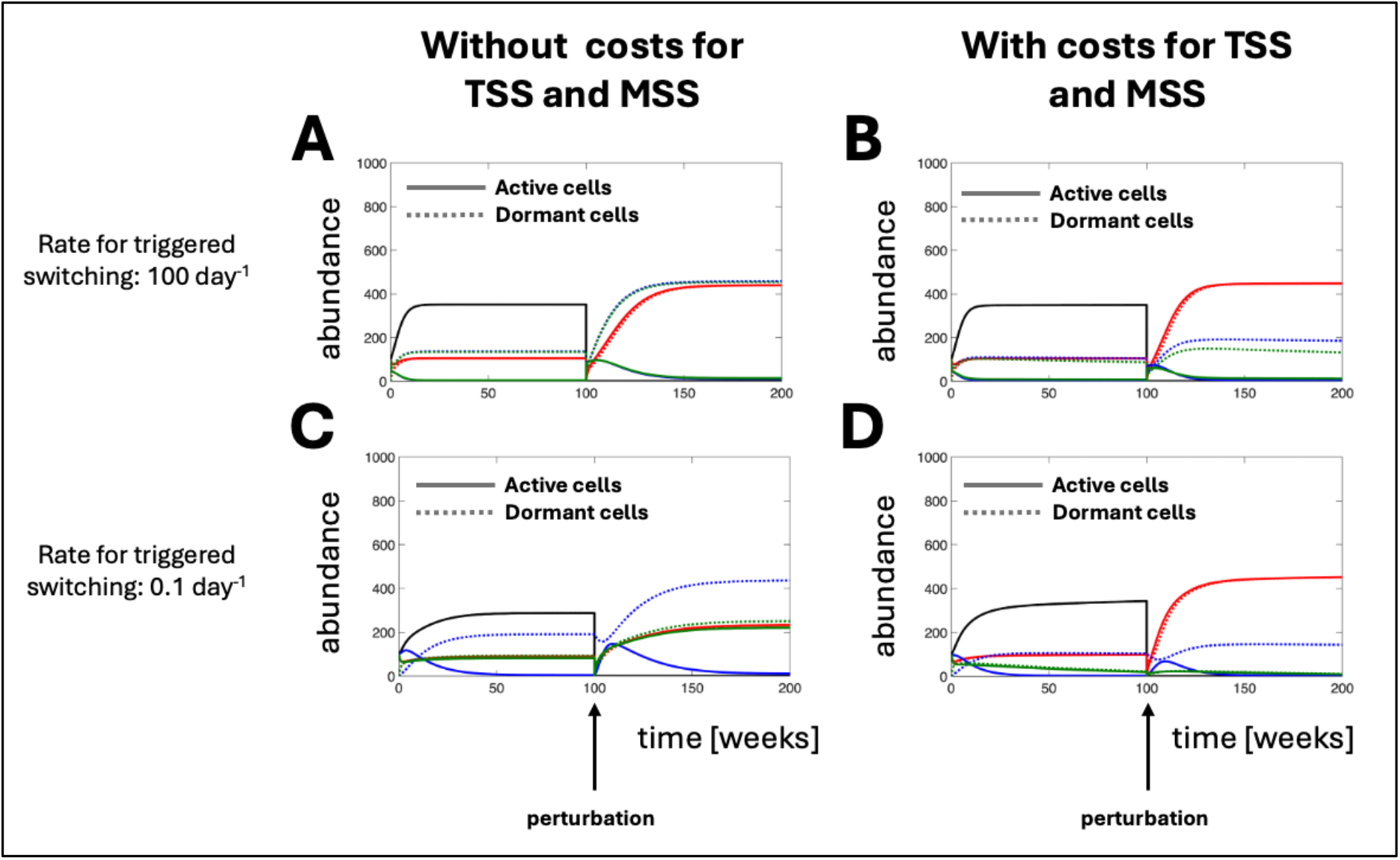
Model dynamics for a perturbed mixed community containing all strategists, comparing binary vs dynamic triggered switching. Similar to figure 3 but comparing different rates s_TD_ for triggered switching. Simulations were initiated with 100 active microbes of each type (and no dormant microbes) and simulated for 200 weeks. The colours show the different strategists: Black: NSS; Red: SSS; Blue: TSS; Green: MSS. Solid lines show active subpopulations; dashed lines indicate dormant subpopulations. **(A)** A community with a high rate for triggered switching in TSS and MSS, such that these strategists are triggered in a binary manner. The community is exposed to a perturbation which removes all active microbes. The perturbation is timed such that no strategist has yet been eliminated. NSS are eliminated by the perturbation; the remaining strategists regenerate their active subpopulations by reactivating surviving dormant cells and compete for the niche resources. After the perturbation, the active community is mainly composed of SSS while TSS and MSS become dormant. **(B)** As in (A) but with a sensing cost, i.e. the growth rate r for TSS and MSS is halved. After the perturbation, the dormant pool of TSS and MSS is smaller compared to (A). **(C)** A community with a lower rate for triggered switching in TSS and MSS, such that these strategists are triggered in a gradual manner. The community is exposed to a perturbation which removes all active microbes. After the perturbation, the pool of active SSS is lower while the pool of active MSS is larger, compared with (A). **(D)** As in (C) but with a sensing cost, i.e. the growth rate r for TSS and MSS is halved. After the perturbation, the dormant pool of MSS is much smaller compared to (C).

